# Controlling stomatal aperture, a potential strategy for managing plant bacterial disease

**DOI:** 10.1101/2022.06.22.497112

**Authors:** Nanami Sakata, Taiki Ino, Chinatsu Hayashi, Takako Ishiga, Yasuhiro Ishiga

**Author notes:** For correspondence: Yasuhiro Ishiga, Address: Faculty of Life and Environmental Sciences, University of Tsukuba, 1-1-1, Tennodai, Tsukuba, Ibaraki 305-8572, Japan.

## Abstract

Bacterial blight of crucifers caused by *Pseudomonas cannabina* pv. *alisalensis* (*Pcal*) inflicts great damage on crucifer production. To explore efficient and sustainable strategies for *Pcal* disease control, we here investigated and screened for amino acids with reduced disease development. We found that exogenous foliar application with multiple amino acids reduced disease symptoms and bacterial populations in cabbage after spray-inoculation, but not syringe-inoculation. These results indicate that these amino acids showed a protective effect before *Pcal* entered plants. Therefore, we observed stomatal responses, which is a main gateway for *Pcal* entry into the apoplast, after amino acid treatments, and found several amino acids induce stomatal closure. Moreover, our findings demonstrated that reducing stomatal aperture width can limit bacterial entry into plants, leading to reduced disease symptoms. Therefore, managing stomatal aperture can be a new powerful strategy for controlling bacterial disease.

## Introduction

Brassicaceae, one of the most important plant families, includes many crop cultivars all around the world. Bacterial blight of crucifers caused by *Pseudomonas cannabina* pv. *alisalensis* (*Pcal*) inflicts great damage on crucifer production including cabbage, Chinese cabbage, and Japanese radish^1,2^. Chemical treatments such as copper fungicides and antibiotics are popular bacterial disease control strategies. However, increasing fungicide resistance is a severe problem, and *Pcal* strains resistant to these chemicals have been reported^3^. Therefore, safer, pollution-free, and sustainable strategies for controlling *Pcal* are valuable for effective bacterial disease control.

Amino acids have been used as water-soluble fertilizers to promote plant growth and improve crop quality^4–7^. In plants, amino acids play essential roles in the regulation of development, growth, and stress response. For example, tryptophan [Trp] is essential for auxin synthesis, which is an important plant growth hormone^8^. Methionine [Met] is an ethylene (ET) precursor, an important hormone in development and stress responses^9^. Isoleucine [Ile] is necessary for jasmonic acid (JA) activation. (+)-7-iso-Jasmonoyl-L-isoleucine is the natural ligand of *Arabidopsis thaliana* COI1-JAZ complexes, and functions as the endogenous bioactive jasmonate^10^. Moreover, previous studies have revealed that amino acids are important in plant disease resistance. For instance, histidine [His] induced resistance in *A. thaliana* and tomato against the fungal pathogen *Botrytis cinerea* and the soilborne bacterial pathogen *Ralstonia solanacearum*^11^. Further, exogenous application with glutamic acid [Glu] enhanced resistance against the fungal pathogen *Magnaporthe oryzae* in rice^12^, *Alternaria alternata* in tomato^13^, and *Colletotrichum higginsianum* and the foliar bacterial pathogen *P. syringae* pv. *tomato* (*Pto*) in *A. thaliana*^14^. Additionally, exogenous isoleucine [Ile] application enhanced *A. thaliana* resistance to *B. cinerea*^15^. Proline [Pro] and alanine [Ala] increased resistance to the Fusarium head blight fungus *Fusarium graminearum*, while cysteine [Cys] aggravated the susceptibility in wheat^16^. These studies indicate that amino acids have potential to control plant disease, but which amino acids are effective in disease control should be carefully considered depending on the plant-pathogen interactions. Moreover, studies on amino acids for bacterial disease control have not been investigated well, and amino acid protective mechanisms against bacterial pathogens have not been elucidated.

The foliar bacterial pathogen *P. syringae* first colonizes leaf surfaces of host plants, enters into plants through natural opening sites, such as stomata and wounds, and then multiplies in the leaf interior^17^. *Pcal* mainly enters the plant through stomata by secreting the phytotoxin coronatine (COR), which functions to reopen stomata^18^, same as the model plant pathogenic bacterium *Pto* DC3000^19^. After entry into the apoplast, bacterial populations often increase greatly, and disease is associated with sites where large internal population sizes have been achieved^20,21^. Additionally, large epiphytic populations are associated with disease severity for foliar bacterial pathogens^22^. However, the reasons why large epiphytic populations are correlated with a great probability of disease occurrence have not been proven. Further, we need to consider the difference in plant-microbe interactions, since *P. syringae* is composed of numerous strains from genetic lineages from both agricultural and environmental habitats^23,24^. For instance, *Pto* DC3000 belongs to phylogroup 1, is a relatively weak epiphyte, but it is a highly aggressive pathogen once inside host tissues^17,24,25,26^. Conversely, *P. syringae* pv. *syringae* (*Psy*) B728a belongs to phylogroup 2, and is a particularly strong epiphytic colonizer ^17,24,25,26^. *Pcal* belongs to phylogroup 5, and its character has been poorly understood^23,24^. We previously found that one of the plant activators, acibenzolar-*S*-methyl (ASM), protects Brassicaceae plants from *Pcal* by activating stomatal-based defense^27–29^. Therefore, it is tempting to speculate that inducing stomatal closure and limiting *Pcal* entry can be an effective way of controlling *Pcal* disease.

We here demonstrated that several amino acids protect plants from *Pcal* infection. Furthermore, these amino acids induce stomatal closure. Moreover, our findings demonstrated that reducing stomatal aperture width can limit bacterial entry into plants, leading to reduced disease symptoms. Therefore, managing stomatal aperture can be a new sustainable strategy for controlling bacterial disease.

## Results

### Amino acids showed a protective effect before *Pcal* entered plants

To investigate the effect of amino acids on disease resistance, we conducted an inoculation assay after amino acid treatments. First, we treated cabbage with 20 amino acids and spray-inoculated with *Pcal* 0 and 24 hours post-treatment (hpt). At 0 hpt, bacterial populations were significantly reduced in cabbage treated with leucine [Leu], Ile, phenylalanine [Phe], valine [Val], asparagine [Asn], Cys, threonine [Thr], Glu, arginine [Arg], His, and lysine [Lys] (Fig. 1a). Disease symptoms were also reduced after these amino acid treatments (Supplementary Fig. S1). At 24 hpt, bacterial populations were significantly reduced in cabbage treated with Leu, Ile, Phe, Pro, Trp, Val, Cys, glycine [Gly], Glu, Arg, His, and Lys (Fig. 1b). Disease symptoms were also reduced after these amino acid treatments (Supplementary Fig. S2). Leu, Ile, Phe, Val, Asn, Cys, Glu, Arg, and Lys reduced disease symptoms (Supplementary Figs. S1, S2) and bacterial populations at both treatments (Figs. 1a,b). Especially, Cys reduced bacterial populations approximately 100 times less than that of the control at 0 hpt (Fig. 1a), and 40 times less at 24 hpt (Fig. 1b). These results indicate that multiple amino acids have the potential to protect plants from bacterial pathogens.

**Figure 1.**
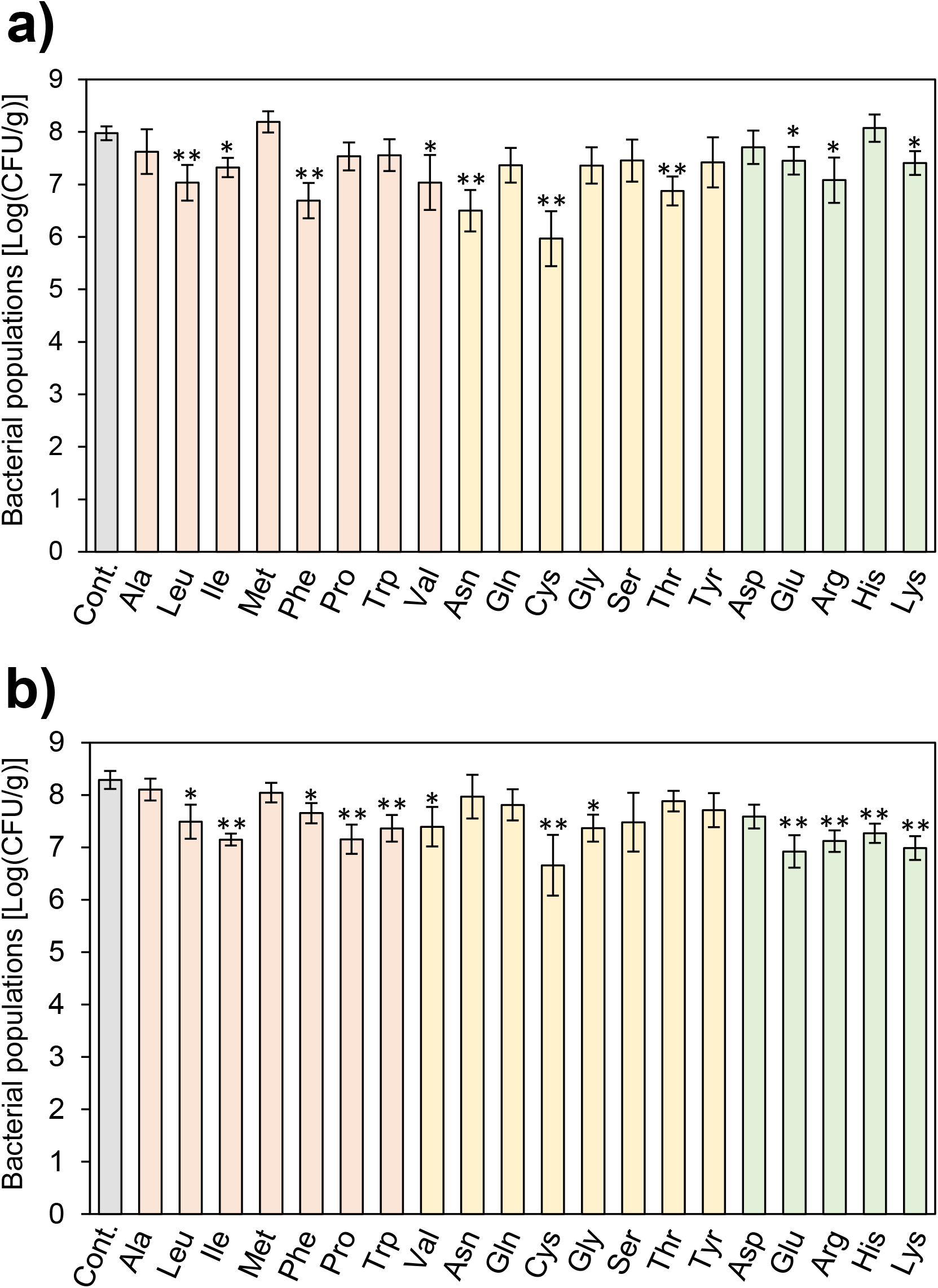
Bacterial populations of amino acids-treated and untreated cabbage leaves after *Pseudomonas cannabina* pv. *alisalnesis* KB211 spray-inoculation. Cabbage plants were spray-inoculated with *Pcal* (5 × 10^7^ CFU/ml) 0 h (a) and 24 h (b) after amino acids spray-treatment (25 mM), including non-polar amino acids (i.e., Ala, Leu, Ile, Met, Phe, Pro, Trp, and Val), polar amino acids (i.e., Asn, Gln, Cys, Gly, Ser, Thr, and Tyr), acidic amino acids (i.e., Asp and Glu), and alkaline amino acids (i.e., Arg, His, and Lys). Water containing 0.025% Tween 20 was used as a control. Vertical bars indicate the standard error for three independent experiments. Asterisks indicate a significant difference from the water treatment control in a *t* test (*<0.05, **<0.01).

To investigate whether these amino acids suppress disease development after *Pcal* enters plants, we conducted syringe-inoculation, which is an inoculation method for directly injecting bacterial suspensions into plants, to bypass stomatal-based defense. Disease symptoms and bacterial populations after amino acid treatments showed no significant differences when syringe-inoculated at 5 × 10^5^ CFU/ml (Fig. 2a; Supplementary Fig. S3). We also conducted syringe inoculation at reduced inoculum levels. There were no significant differences in bacterial populations at the 5 × 10^4^ CFU/ml and 5 × 10^3^ CFU/ml inoculum levels after amino acid treatments (Figs. 2b,c). Disease symptoms were also the same as the control (Supplementary Figs. S4, S5). Together, these results indicate that amino acids showed a protective effect before *Pcal* entered plants.

**Figure 2.**
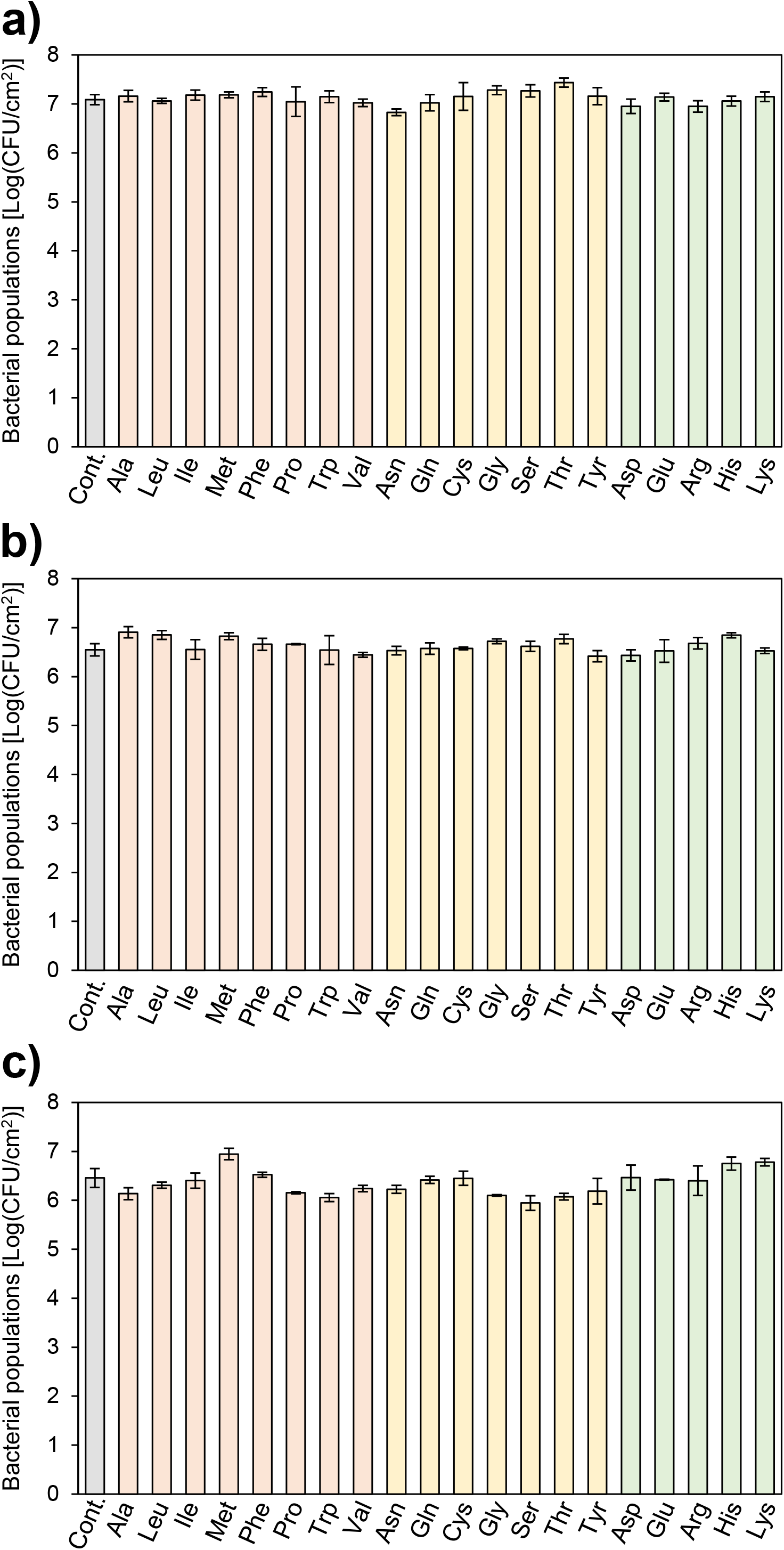
Bacterial populations of amino acids-treated and untreated cabbage leaves after *Pseudomonas cannabina* pv. *alisalnesis* KB211 syringe-inoculation. Cabbage plants were syringe-inoculated with 5 × 10^5^ CFU/ml (a), 5 × 10^4^ CFU/ml (b), and 5 × 10^3^ CFU/ml (c) of inoculum 24 h after amino acids spray-treatment (25 mM), including non-polar amino acids (i.e., Ala, Leu, Ile, Met, Phe, Pro, Trp, and Val), polar amino acids (i.e., Asn, Gln, Cys, Gly, Ser, Thr, and Tyr), acidic amino acids (i.e., Asp and Glu), and alkaline amino acids (i.e., Arg, His, and Lys). Water was used as a control. Vertical bars indicate the standard error for three independent experiments.

### Several amino acids reduced stomatal aperture width

Amino acids showed a protective effect after spray-inoculation but not syringe-inoculation (Figs. 1, 2), so we observed stomatal response, which is a main gateway for *Pcal* entry into the apoplast, after amino acid treatments. Several amino acids reduced the stomatal aperture width (Fig. 3). Especially, stomatal aperture width after Cys treatments were significantly reduced to around 0.71 μm (Fig. 3). Leu, Trp, Gly, Glu, and Lys also induced greater stomatal closure compared to other amino acids (Fig. 3). Amino acids (Ala, Met, Gln, Ser, Tyr, and Asp), which did not eilicit significantly different disease severity from the control, showed no significant or relatively less reduction of stomatal aperture width. Therefore, we hypothesized that most of the amino acids showed a protective effect by inducing stomatal closure and limiting bacterial entry.

**Figure 3.**
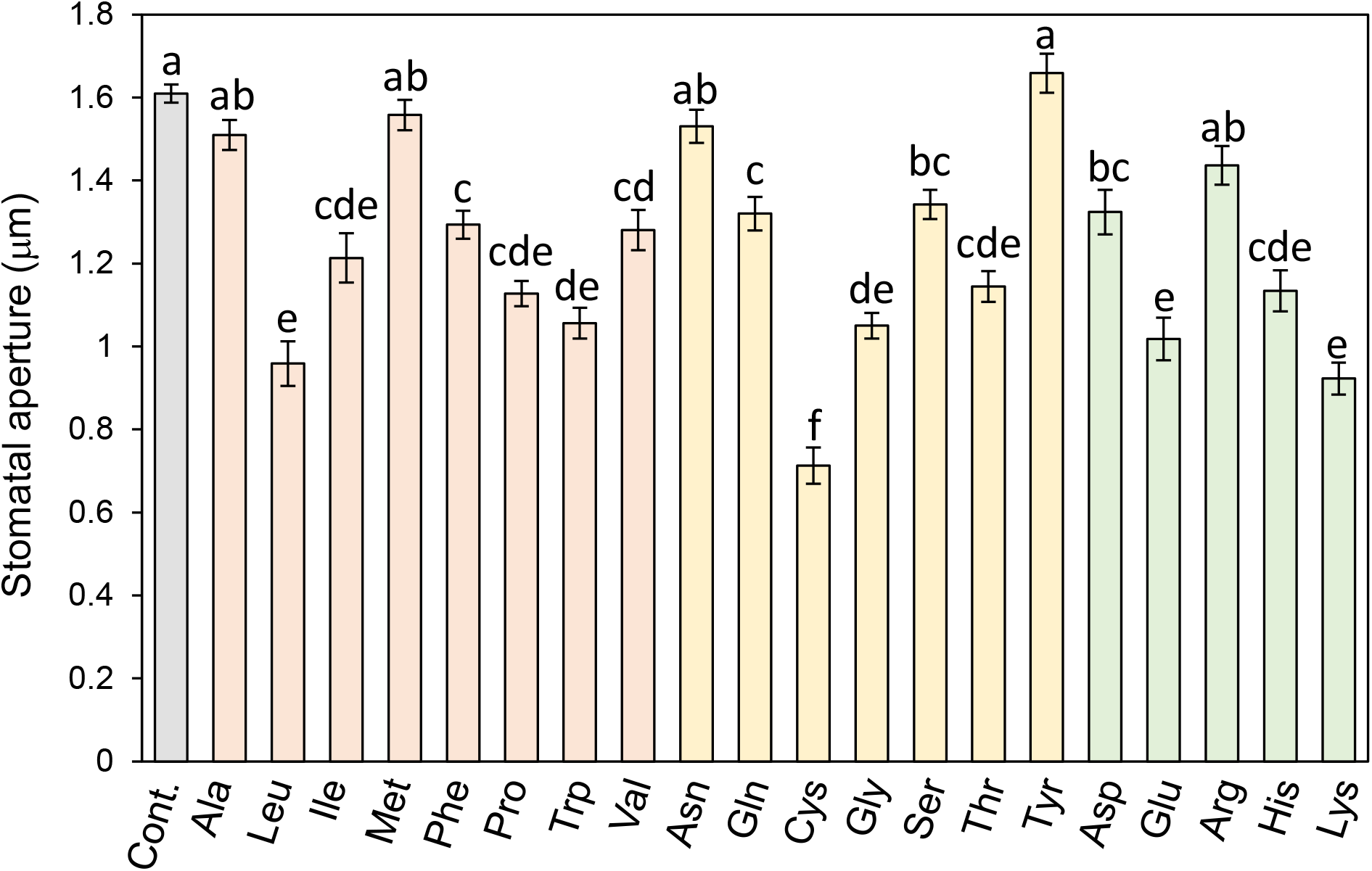
Stomatal response after amino acid treatments. Cabbage leaves were spray-treated with amino acids (25 mM), including non-polar amino acids (i.e., Ala, Leu, Ile, Met, Phe, Pro, Trp, and Val), polar amino acids (i.e., Asn, Gln, Cys, Gly, Ser, Thr, and Tyr), acidic amino acids (i.e., Asp and Glu), and alkaline amino acids (i.e., Arg, His, and Lys). The microscopy leaf images were taken 4 h after treatment using a Nikon optical microscope (Eclipse 80i). The stomatal aperture width (μm) was analyzed by Image J using at least 114 stomata. Vertical bars indicate the standard error for three independent experiments. Different letters indicate a significant difference among treatments based on Tukey’s honestly significant difference test (*p* <0.05).

### Initial bacterial entry correlated with disease symptoms

To test our hypothesis described above, we first investigated whether stomatal aperture width and bacterial entry were correlated or not. To manage stomatal aperture width stepwise, we controlled relative humidity (RH) and kept plants under these conditions. Stomatal aperture width was greater as the RH increased (Fig. 4a). We next investigated bacterial entry at each experimental condition. Bacterial entry was also greater as the RH increased (Fig. 4b). Correlation analysis demonstrated a strong positive correlation between stomatal aperture width and bacterial entry (R^2^=0.95) (Fig. 4c). These results indicate that bacterial entry and stomatal aperture width were correlated.

**Figure 4.**
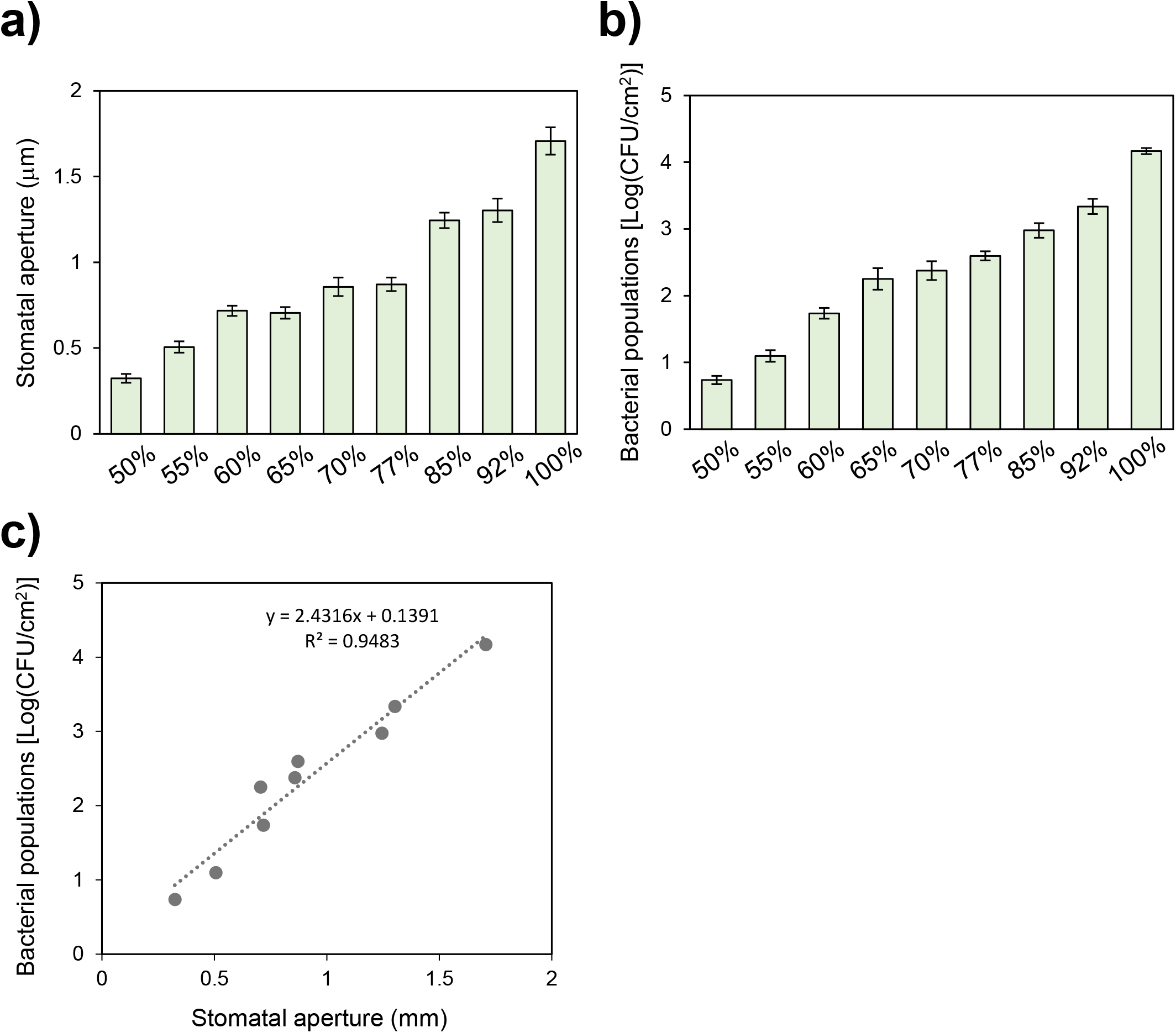
Stomatal response and bacterial entry at different humidity levels. Stomatal aperture width (μm) (a) and bacterial entry (b) at different humidity levels (50%, 55%, 60%, 65%, 70%, 77%, 85%, 92%, and 100% RH). Cabbage plants were placed in the chamber and maintained at different humidity levels for 24-48 h. Cabbage leaves were spray-inoculated with *Pcal* (1 × 10^8^ CFU/ml). The microscopy leaf images were taken 4 h after inoculation using a Nikon optical microscope (Eclipse 80i). The stomatal aperture width (μm) was analyzed by Image J using at least 85 stomata. Bacterial populations were measured at 4 hpi. Vertical bars indicate the standard error for three independent experiments. (c) Correlation analysis between stomatal width and bacterial entry.

Next, to investigate whether bacterial entry correlated with disease symptom development, we syringe-inoculated *Pcal* into cabbage at different inoculum levels. Initial bacterial entry was correlated with disease symptoms (Figs. 5a,b). Necrosis symptoms were observed when we inoculated at 5 × 10^5^ CFU/ml, but were not observed at the 5 × 10^4^ CFU/ml inoculum level (Figs. 5a,b). Moreover, chlorosis was reduced more than two times at the 5 × 10^3^ CFU/ml inoculum level compared to at the 5 × 10^4^ CFU/ml inoculum level, and few chlorosis symptoms were observed at the 5 × 10^2^ CFU/ml inoculum level (Figs. 5a,b). Bacterial populations also increased in proportion to the initial inoculum level (Fig. 5c). These results suggest that the initial bacterial entry threshold for necrosis formation is approximately between 5 × 10^2^ to 5 × 10^3^ CFU/cm^2^, and that for chlorosis formation is approximately between 5 to 50 CFU/cm^2^ (Figs. 5a-c). These results highly support that bacterial entry is correlated with disease development. Taken together, our results suggest that controlling stomatal aperture width can limit bacterial entry into plants, leading to the reduction of disease symptoms.

**Figure 5.**
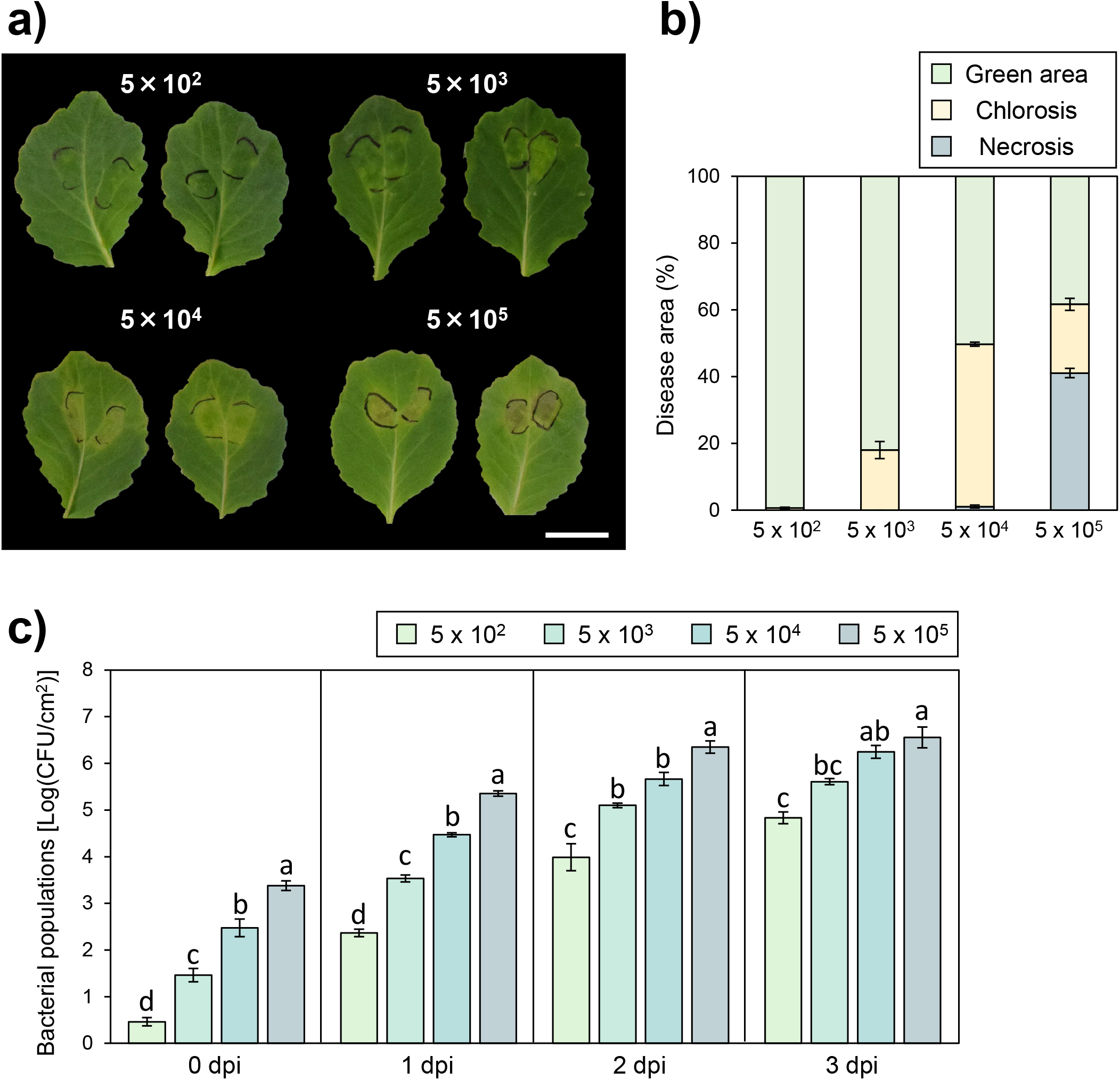
Necrosis and chlorosis area and bacterial populations in cabbage after *Pcal* inoculation. Disease symptoms (a) and percentage of necrosis and chlorosis areas (b) after *Pcal* inoculation at 3 dpi. Bacterial populations (c) after *Pcal* inoculation at 0, 1, 2, and 3 dpi. Cabbage plants were syringe-inoculated with 5x 10^2^ CFU/ml, 5x 10^3^ CFU/ml, 5x 10^4^ CFU/ml, and 5x 10^5^ CFU/ml. Scale bars show 2 cm. Necrosis and chlorosis areas were detected by the ImageJ-based PIDIQ method as described in the methods. Vertical bars indicate the standard error for three independent experiments. Different letters indicate a significant difference among inoculum levels based on Tukey’s honestly significant difference test (*p* <0.05).

### Cys, Glu, and Lys induced stomatal closure and reduced bacterial entry

Our results suggest that reducing stomatal aperture width leads to limiting bacterial entry and disease symptoms (Figs. 4, 5). Therefore, we next investigated whether amino acids, which showed a protective effect, induced stomatal closure and reduce bacterial entry. We tested amino acids (Leu, Trp, Cys, Gly, Glu, and Lys), which showed less than 1.1 μm stomatal aperture after treatment without inoculation, and Ala as a negative control. Stomatal aperture width was particularly reduced to less than 0.8 μm after Trp, Cys, Gly, Glu, and Lys treatment (Fig. 6a). Bacterial entry was significantly reduced after Cys, Glu, and Lys treatment (Fig. 6b). Cys closed stomata to less than 0.45 μm and suppressed bacterial entry to around 5 × 10^2^ CFU/cm^2^ (Figs. 6a,b). These results suggest that Cys, Glu, and Lys induced stomatal closure and reduced bacterial entry, leading to a reduction in disease severity.

**Figure 6.**
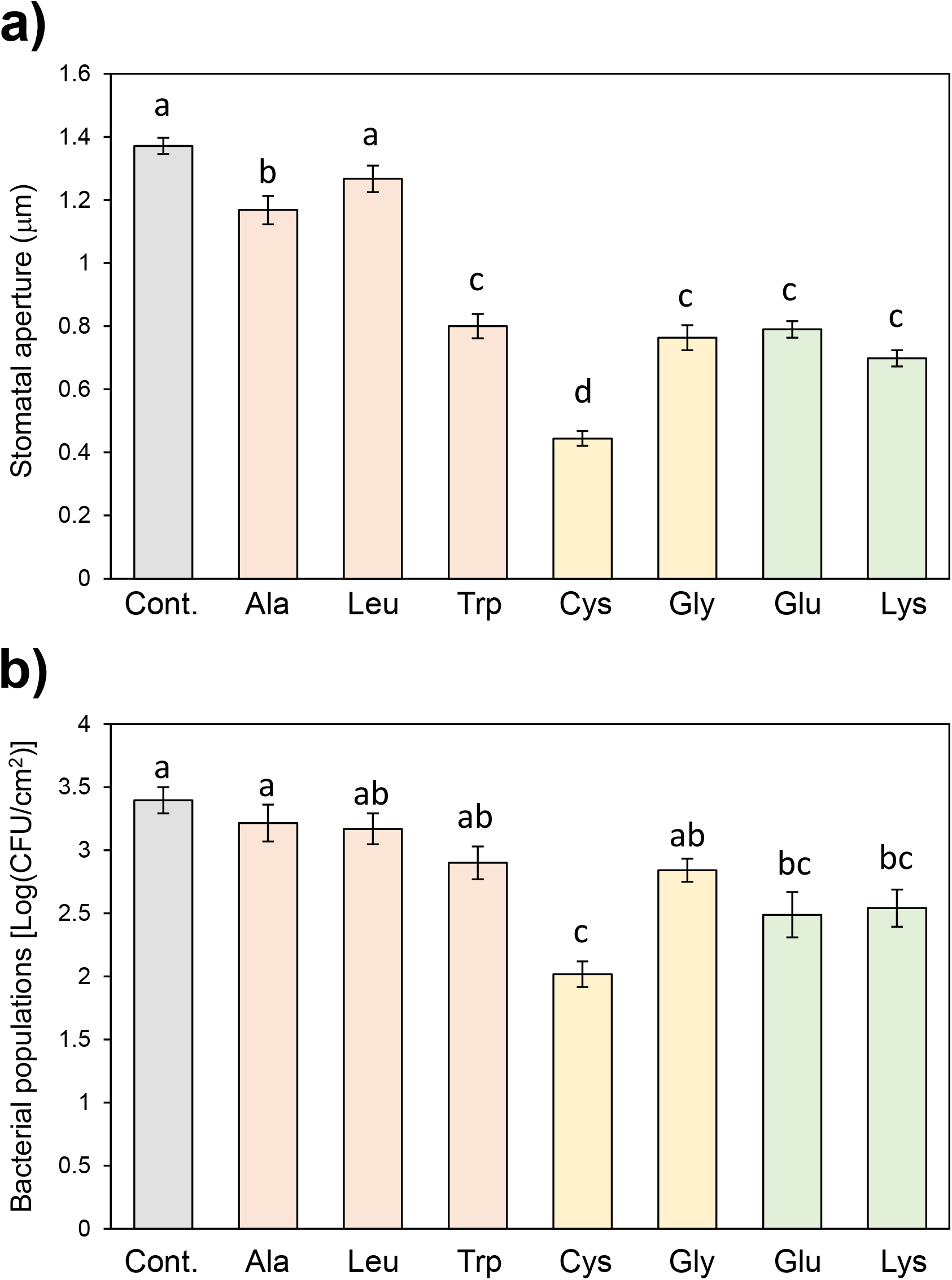
Stomatal responses of amino acids-treated and untreated leaves after *Pcal* inoculation. Stomatal aperture width (μm) (a) and bacterial entry (b) after aino acids (Ala, Leu, Trp, Cys, Gly, Glu, and Lys) treatments and *Pcal* inoculation. Cabbage leaves were spray-treated with amino acids (25 mM) and leaves were spray-inoculated with *Pcal* (1 × 10^8^ CFU/ml) at 4 hpt. The microscopy leaf images were taken 4 h after inoculation using a Nikon optical microscope (Eclipse 80i). The stomatal aperture width (μm) was analyzed by Image J using at least 310 stomata. Bacterial populations were measured at 4 hpi. Vertical bars indicate the standard error for three independent experiments. Different letters indicate a significant difference among treatments based on Tukey’s honestly significant difference test (*p* <0.05).

## Discussions

Previous studies revealed that several amino acids induced plant resistance against plant pathogens. However, studies on amino acids for bacterial disease control have not been investigated well, and the protective mechanisms of amino acids against bacterial pathogens have not been elucidated. Thus, we investigated if amino acids treatment resulted in reduced disease development. Several amino acids reduced *Pcal* disease and induced stomatal closure. We also demonstrated that reducing stomatal aperture width leads to a decrease in bacterial entry, resulting in a decrease in disease symptoms. Our results shed light on the importance of managing the stomatal aperture, which is a gateway for bacterial entry, for controlling bacterial pathogens.

Exogenous foliar application with multiple amino acids reduced disease symptoms and bacterial populations in cabbage spray-inoculated with *Pcal* (Figs. 1; Supplementary Figs. S1, S2). However, no amino acids reduced disease symptoms and bacterial populations in syringe-inoculated cabbage (Figs. 2; Supplementary Figs. S3, S4, S5). These results indicate that amino acids had a protective effect before *Pcal* entered plants. Several studies showed amino acids contribute to plant resistance by inducing plant phytohormones, such as salicylic acid (SA), JA, and ET. For instance, Glu-induced blast resistance in rice partly depends on the SA-mediated signaling pathway^12^. Ile enhanced resistance against *B. cinerea* via the JA-mediated signaling pathway^15^. His induced resistance to *R. solanacearum* partly through activation of the ET-mediated signaling pathway^11^. Thus, our results provide new insight into amino acid protection mechanisms against bacterial disease, and shed light on the importance of protecting plants from bacteria before bacterial invasion.

Multiple amino acids induced stomatal closure (Fig. 3). Plant glutamate receptor homologs (GLRs) are involved in various Ca^2+^-mediated development regulations and physiological responses, including stomatal closure^30^. Additionally, amino acids Ala, Asn, Cys, Gly, Glu, and Ser act as agonists for plant GLRs^31–34^. Cys induced the greatest stomatal closure regardless of *Pcal* inoculation (Figs. 3, 6). Cys triggers stomatal closure by inducing abscisic acid (ABA) biosynthesis^35^. Moreover, γ-aminobutyric acid (GABA) acts as an endogenous plant signaling molecule and modulates stomatal opening to enhance plant water use efficacy and drought resilience^36^. Therefore, it is tempting to speculate that amino acids have the potential to modulate stomatal closing.

If initial bacterial entry was related to visible lesion formation, we would expect disease severity. Based on our study, it seems the initial bacterial entry threshold for necrosis formation is approximately between 5 × 10^2^ to 5 × 10^3^ CFU/cm^2^, and that for chlorosis formation is approximately between 5 to 50 CFU/cm^2^ (Figs. 5). Our results also showed that stomatal aperture width correlated with bacterial entry numbers (Figs. 4). Therefore, we propose that controlling stomatal aperture width can limit bacterial entry into plants, leading to disease symptom reduction. The concept of “infection threshold” was used to predict disease incidence at the field level^37,38^. Bacterial populations often increase greatly, and disease is associated with sites where large internal population sizes have been achieved^20,21^. Moreover, large epiphytic populations are associated with a diease onset time, and with increased disease severity for foliar bacterial pathogens^22^. It is thought that the major role of epiphytic pathogen populations in disease development is probably to increase endophytic population sizes^22^. Our results clearly showed the quantitative relationship among stomatal aperture width, bacterial entry, and disease development in *Pcal*-cabbage interactions, and strongly support this idea.

Cys, Glu, and Lys treatments showed significantly reduced stomatal closure and bacterial entry (Figs. 6). These three amino acids reduced disease symptoms and bacterial growth both at 0 and 24 hpt (Figs. 1). These results indicate that Cys, Glu, and Lys treatments have the potential to be effective bacterial control strategies. Conversely, although Leu reduced stomatal aperture width without *Pcal* inoculation (Fig. 3), no significant differences in stomatal aperture width with *Pcal* inoculation were observed (Figs. 6). Moreover, Trp and Gly reduced stomatal aperture width during *Pcal* infection, but these amino acids had less effect on bacterial entry (Figs. 6). Indeed, based on our model in Fig. 4c, if stomatal aperture width is less than 0.5 μm, bacterial entry is around 1.4 CFU/cm^2^ or less. However, in Fig. 6, the bacterial entry was around 2 CFU/cm^2^, when the stomatal aperture width was 0.44 μm (Cys treatment). These results indicate that the actual bacterial entry in Fig. 6 was greater than expected based on our model in Fig. 4c. There are several possible explanations for these gaps. First, we must consider the effect of amino acids on bacterial virulence factors and plant resistance. *Pcal* reopens stomata by secreting COR, which is one of the main virulence factors^18^. Therefore, it can be considered that some amino acids suppress bacterial virulence factors, which function for suppressing stomatal-based defense, such as COR and type three effectors (TTEs). Further, there is a great possibility that the greater growth than we expected occurred by increased humidity conditions, since we conducted the experiment at 100% RH in Figs. 6. In the field, phyllosphere bacterial disease outbreaks typically occur after rainfall and high humidity conditions^39^. Moreover, *Pto* DC3000 effectors, such as HopM1, are induced in the aqueous apoplast, and are sufficient to transform non-pathogenic *P. syringae* strains into virulence pathogens in immunodeficient *A. thaliana* under high humidity^40^. An aqueous apoplast could potentially facilitate bacterial nutrient uptake, promote bacterial spread, and/or affect apoplastic host defense responses^40^. Although we cannot exclude the possibility of the other effects of amino acids, our results suggest that amino acids suppressed *Pcal* disease by limiting bacterial entry through inducing stomatal closure.

Our studies shed light on the importance of stomatal management, which is a gateway for bacterial entry. Stomata have an important role in exchanging gas and environmental adaptation^41^. Importantly, it is not necessary to close stomata completely to reduce disease symptoms. Our results suggest that reducing stomatal aperture width moderately is enough to limit bacterial entry and disease symptom development. Therefore, managing stomatal aperture moderately can be a new powerful strategy for controlling bacterial diseases.

## Materials and methods

### Plant materials and growth conditions

Cabbage (*Brassica oleracea* var. *capitate*) cv. Kinkei 201 was used for *Pcal* virulence assays. Cabbage was grown from seed at 23-25°C with a light intensity of 200 μEm^−2^s^−1^ and a 16-h light/8-h dark photoperiod. Cabbage plants were used for spray- and syringe-inoculation assays around two weeks after germination.

### Application of exogenous amino acids

L-Ala (Tokyo Chemical Industry, Tokyo, Japan), L-leu (Nacalai tesque, INC., Kyoto, Japan), L-Ile (Tokyo Chemical Industry), L-Met (Tokyo Chemical Industry), L-Phe (Tokyo Chemical Industry), L-Pro (Tokyo Chemical Industry), L-Trp (FUJIFILM Wako Pure Chemical Corporation, Osaka, Japan), L-Val (Tokyo Chemical Industry), L-Asn (Tokyo Chemical Industry), L-Gln (Tokyo Chemical Industry), L-Cys (Tokyo Chemical Industry), L-Ser (FUJIFILM Wako Pure Chemical Corporation), L-Thr (Tokyo Chemical Industry), L-Tyr (Tokyo Chemical Industry), L-Asp (Tokyo Chemical Industry), L-Glu (FUJIFILM Wako Pure Chemical Corporation), L-Arg (Tokyo Chemical Industry), L-His (Tokyo Chemical Industry), and L-Lys (Nacalai tesque, INC.) were dissolved in sterile distilled H_2_O containing 0.025% (v/v) Tween 20 (FUJIFILM Wako Pure Chemical Corporation) solution to prepare amino acid solutions at 25 mM. To evaluate the amino acids effect on disease control, cabbage leaves were spray-treated with amino acids (25 mM) at 0 and 24 hpt. Plants were spray-treated with water containing 0.025% Tween 20 as controls.

### Bacterial strains, plasmids, and growth conditions

Pathogenic *Pseudomonas cannabina* pv. *alisalensis* strain KB211^42^ was kindly provided by the Nagano Vegetable and Ornamental Crop Experiment Station, Nagano, Japan. *Pcal* wild-type was grown on King’s B ^43^ medium at 28°C. Before *Pcal* inoculation, bacteria were suspended in sterile distilled H_2_O, and the bacterial cell densities at 600 nm (OD_600_) were measured using a Biowave CO8000 Cell Density Meter (Funakoshi, Tokyo, Japan).

### Bacterial inoculation methods

To assay for cabbage disease, spray-inoculation was conducted by spraying seedlings with bacterial suspensions (5 × 10^7^ colony forming units: CFU/ml) containing 0.025% Silwet L-77 (OSI Specialities, Danbury, CT, USA) until runoff. The seedlings were then incubated in growth chambers at 85-95% RH for the rest of the experimental period. Leaves were removed and photographed at 5 dpi. For syringe-inoculation, plants were syringe-inoculated with bacterial suspensions (5 × 10^2^, 5 × 10^3^, 5 × 10^4^, and 5 × 10^5^ CFU/ml) with a 1-ml blunt syringe into leaves. The plants were then incubated at 70-80% RH for the rest of the experimental period. Leaves were removed and photographed at 3 dpi.

To assess bacterial growth in cabbage, the internal bacterial populations were measured after spray-inoculation. Inoculated seedlings were collected, and a total of two inoculated leaves were measured to quantify the bacterial populations. The leaves were surface sterilized with 10% H_2_O_2_ for 3 minutes. After washing three times with sterile distilled water, the leaves were homogenized in sterile distilled water, and diluted samples were plated onto solid KB agar medium. For syringe-inoculation, to assess bacterial growth in cabbage, leaf discs were harvested using a 3.5 mm-diameter cork borer from syringe-infiltrated leaf zones. The plant samples were homogenized in sterile distilled water, and diluted samples were plated onto solid KB agar medium. Two or three days after dilution sample plating, the bacterial colony forming units (CFUs) were counted and normalized as CFU per gram or CFU per cm^2^, using the total leaf weight or leaf square centimeters. The bacterial populations at 0 dpi were estimated using leaves harvested 1-hour post-inoculation (hpi) without surface-sterilization. The bacterial populations were evaluated in at least three independent experiments.

### Bacterial entry assay

To assess bacterial entry into the plant apoplast through stomata, the internal bacterial population was measured after spray-inoculation. Cabbage were spray-inoculated with bacterial suspensions (1 × 10^8^ CFU/ml) containing 0.025% Silwet L-77 (OSI Specialities) until runoff. After 4 hpi, inoculated seedlings were collected, and surface sterilized with 10% H_2_O_2_ for 3 minutes. After washing three times with sterile distilled water, leaf discs were harvested using a 3.5 mm-diameter cork-borer. The plant samples were homogenized in sterile distilled water, and diluted samples were plated onto solid KB agar medium. Two or three days after dilution sample plating, the bacterial CFUs were counted and normalized as CFU per cm^2^. The bacterial populations were evaluated in at least three independent experiments.

### Stomatal assay

A modified method was used to assess stomatal response as described previously^44^. Briefly, cabbage plants were grown for 2 weeks after sowing. For the stomatal assay in response to 20 amino acids, cabbage leaves were cut and floated on the amino acid suspensions, respectively. Amino acid treated, or water mock-treated leaves were directly imaged at 4 hpt using a Nikon optical microscope (Eclipse 80i). The aperture width of at least 114 stomata was measured at 4 hpt.

For the stomatal assay in response to different humidity conditions, plants were maintained at 50%, 55%, 60%, 65%, 70%, 77%, 85%, 92%, and 100% RH for 24-48 h. The treated leaves were then spray-inoculated with *Pcal* (1 × 10^8^ CFU/ml). Spray-inoculated or water mock-inoculated cabbage leaves were directly imaged at 4 hpi using a Nikon optical microscope (Eclipse 80i). The aperture width of at least 85 stomata was measured.

For the stomatal assay in response to select amino acids (Ala, Leu, Trp, Cys, Gly, Glu, and Lys) and *Pcal* inoculation, cabbage plants were spray treated with these amino acids until runoff. The plants were then incubated at 100% RH for 24 h. The treated leaves were then spray-inoculated with *Pcal* (1 × 10^8^ CFU/ml) at 24 hpt. Spray-inoculated or water mock-inoculated cabbage leaves were directly imaged at 4 hpi using a Nikon optical microscope (Eclipse 80i). The aperture width of at least 310 stomata was measured.

The stomatal aperture width was measured using the ImageJ software. The average and SE for the stomatal aperture width were calculated.

### Measurement of disease area

To measure healthy and lesion-bearing areas of cabbage infected with *Pcal*, ImageJ Ver. 1.53e (Waye Rasband and contributors, National Institutes of Health, USA)-based macro plant immunity and disease image-based quantification (PIDIQ) was used^45–47^. The ‘Threshold Color’ values for necrosis and chlorosis were adjusted as described in the supplemental data (Supplementary ImageJ macro 1 and 2), respectively. The workflow and images for lesions areas detected by the ImageJ-based PIDIQ method are shown in Supplementary Figs. 6.

## Statistical Analysis

All data are expressed as the mean with SE. All statistical analyses were performed using EZR^48^, a graphical user interface for R (version 3.6.3; R Foundation for Statistical Computing, Vienna, Austria). Tukey’s honestly significant difference (HSD) test was used to analyze data. Differences of *p* < 0.05 were considered statistically significant.

## Data availability

The data presented in this study are openly available in the supplementary materials here.

## Acknowledgments

We thank Dr. Christina Baker for editing the manuscript. We thank Dr. Hidenori Matsui of Okayama University for the technical assistance. *Pcal* was kindly given from the Nagano vegetable and ornamental crops experiment station, Nagano, Japan. This work was supported, in part, by the Japan Society for the Promotion of Science (JSPS), Grant Number: 19K06045 (Y.I.), and by the JSPS, Grant Number: 21J10765 (N.S.).

## Author contributions

N.S. and Y.I. designed the experiments; N.S., T.I., C.H., T.I., and Y.I. performed the experiments; N.S. and Y.I. wrote the manuscript.

## Supplementary information

**Supplementary Fig. S1. Disease symptoms of amino acids-treated and untreated cabbage leaves after *Pseudomonas cannabina* pv. *alisalnesis* KB211 spray-inoculation**. Cabbage plants were spray-inoculated with *Pcal* (5 × 10^7^ CFU/ml) 0 h after amino acids spray-treatment (25 mM), including non-polar amino acids (i.e., Ala, Leu, Ile, Met, Phe, Pro, Trp, and Val), polar amino acids (i.e., Asn, Gln, Cys, Gly, Ser, Thr, and Tyr), acidic amino acids (i.e., Asp and Glu), and alkaline amino acids (i.e., Arg, His, and Lys). Water containing 0.025% Tween 20 was used as a control. The leaves were photographed at 5 dpi. Scale bars show 2 cm.

**Supplementary Fig. S2. Disease symptoms of amino acids-treated and untreated cabbage leaves after *Pseudomonas cannabina* pv. *alisalnesis* KB211 spray-inoculation**. Cabbage plants were spray-inoculated with *Pcal* (5 × 10^7^ CFU/ml) 24 h after amino acids spray-treatment (25 mM), including non-polar amino acids (i.e., Ala, Leu, Ile, Met, Phe, Pro, Trp, and Val), polar amino acids (i.e., Asn, Gln, Cys, Gly, Ser, Thr, and Tyr), acidic amino acids (i.e., Asp and Glu), and alkaline amino acids (i.e., Arg, His, and Lys). Water containing 0.025% Tween 20 was used as a control. The leaves were photographed at 5 dpi. Scale bars show 2 cm.

**Supplementary Fig. S3. Disease symptoms of amino acids-treated and untreated cabbage leaves after *Pseudomonas cannabina* pv. *alisalnesis* KB211 syringe-inoculation**. Cabbage plants were syringe-inoculated with 5x 10^5^ CFU/ml of inoculum 24 h after amino acids spray-treatment (25 mM), including non-polar amino acids (i.e., Ala, Leu, Ile, Met, Phe, Pro, Trp, and Val), polar amino acids (i.e., Asn, Gln, Cys, Gly, Ser, Thr, and Tyr), acidic amino acids (i.e., Asp and Glu), and alkaline amino acids (i.e., Arg, His, and Lys). Water was used as a control. The leaves were photographed at 5 dpi. Scale bars show 2 cm.

**Supplementary Fig. S4. Disease symptoms of amino acids-treated and untreated cabbage leaves after *Pseudomonas cannabina* pv. *alisalnesis* KB211 syringe-inoculation**. Cabbage plants were syringe-inoculated with 5x 10^4^ CFU/ml of inoculum 24 h after amino acids spray-treatment (25 mM), including non-polar amino acids (i.e., Ala, Leu, Ile, Met, Phe, Pro, Trp, and Val), polar amino acids (i.e., Asn, Gln, Cys, Gly, Ser, Thr, and Tyr), acidic amino acids (i.e., Asp and Glu), and alkaline amino acids (i.e., Arg, His, and Lys). Water was used as a control. The leaves were photographed at 5 dpi. Scale bars show 2 cm.

**Supplementary Fig. S5. Disease symptoms of amino acids-treated and untreated cabbage leaves after *Pseudomonas cannabina* pv. *alisalnesis* KB211 syringe-inoculation**. Cabbage plants were syringe-inoculated with 5x 10^3^ CFU/ml of inoculum 24 h after amino acids spray-treatment (25 mM), including non-polar amino acids (i.e., Ala, Leu, Ile, Met, Phe, Pro, Trp, and Val), polar amino acids (i.e., Asn, Gln, Cys, Gly, Ser, Thr, and Tyr), acidic amino acids (i.e., Asp and Glu), and alkaline amino acids (i.e., Arg, His, and Lys). Water was used as a control. The leaves were photographed at 5 dpi. Scale bars show 2 cm.

**Supplementary Fig. S6. Necrosis and chlorosis area of cabbage plants after *Pcal* inoculation**. (a) Workflow of the PIDIQ method. The area that has been circled with a white dotted line was selected as an inoculated area by using Adobe Photoshop 2022. (b) Necrosis areas detected by the ImageJ-based PIDIQ method as described in the methods. Necrosis and other areas are colored in blue and red, respectively. (c) Chlorosis areas detected by the ImageJ-based PIDIQ method as described in the methods. Chlorosis and other areas are colored in blue and red, respectively.

**Supplementary ImageJ macro 1 & 2**. The ‘Threshold Color’ values for necrosis and chlorosis.

**Supplementary data sheets**. The data presented in this study.

