## Supplemental Figures for "Controlling stomatal aperture, a potential strategy for managing plant bacterial disease"

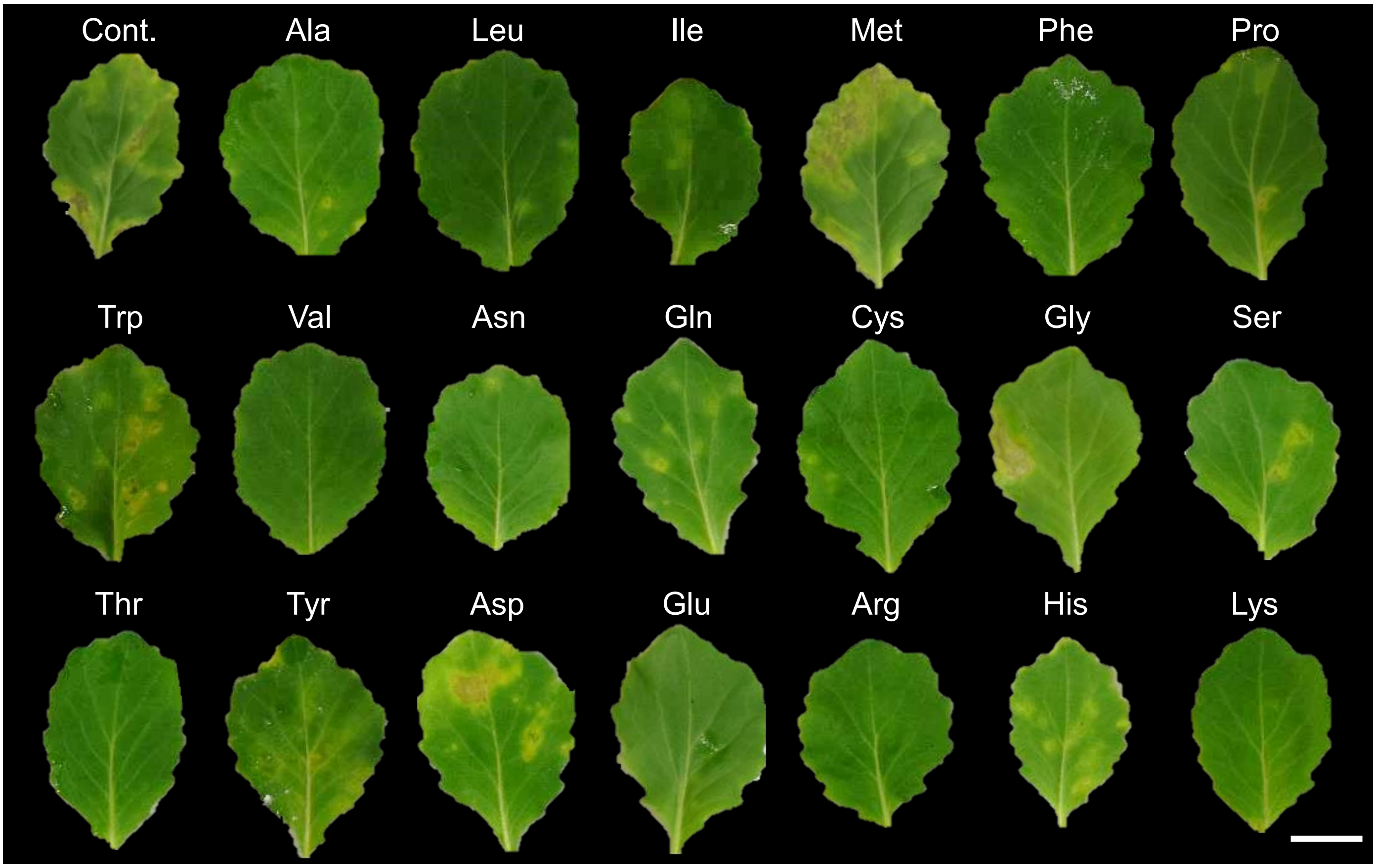


**Supplementary Fig. S1. Disease symptoms of amino acids-treated and untreated cabbage leaves after *Pseudomonas cannabina* pv. *alisalnesis* KB211 spray-inoculation.** Cabbage plants were spray-inoculated with *Pcal* (5 x $10^7$ CFU/ml) 0 h after amino acids spray-treatment (25 mM), including non-polar amino acids (i.e., Ala, Leu, Ile, Met, Phe, Pro, Trp, and Val), polar amino acids (i.e., Asn, Gln, Cys, Gly, Ser, Thr, and Tyr), acidic amino acids (i.e., Asp and Glu), and alkaline amino acids (i.e., Arg, His, and Lys). Water containing 0.025% Tween 20 was used as a control. The leaves were photographed at 5 dpi. Scale bars show 2 cm.

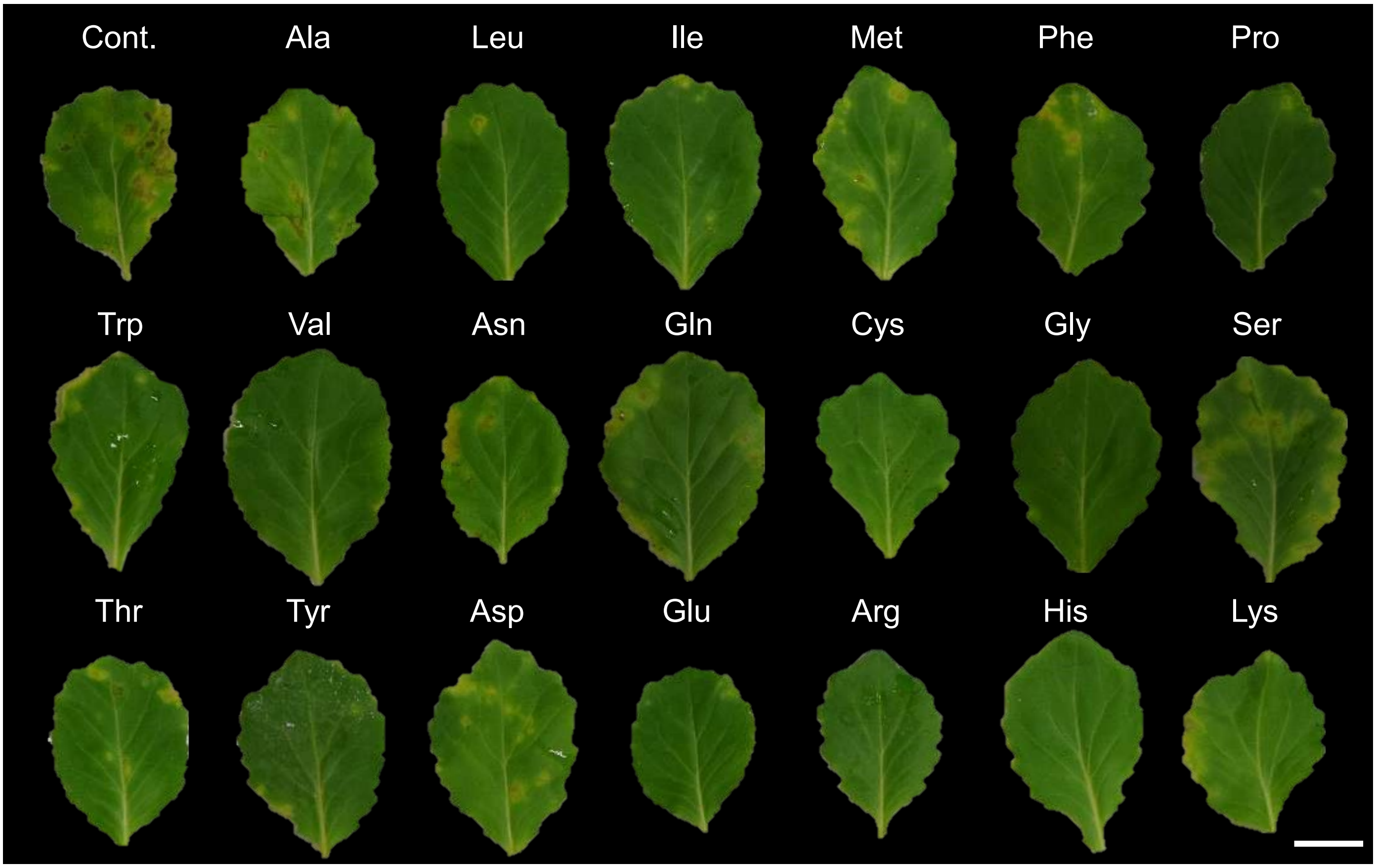


**Supplementary Fig. S2**. **Disease symptoms of amino acids-treated and untreated cabbage leaves after *Pseudomonas cannabina* pv. *alisalnesis* KB211 spray-inoculation.** Cabbage plants were spray-inoculated with *Pcal* (5 x $10^7$ CFU/ml) 24 h after amino acids spray-treatment (25 mM), including non-polar amino acids (i.e., Ala, Leu, Ile, Met, Phe, Pro, Trp, and Val), polar amino acids (i.e., Asn, Gln, Cys, Gly, Ser, Thr, and Tyr), acidic amino acids (i.e., Asp and Glu), and alkaline amino acids (i.e., Arg, His, and Lys). Water containing 0.025% Tween 20 was used as a control. The leaves were photographed at 5 dpi. Scale bars show 2 cm.

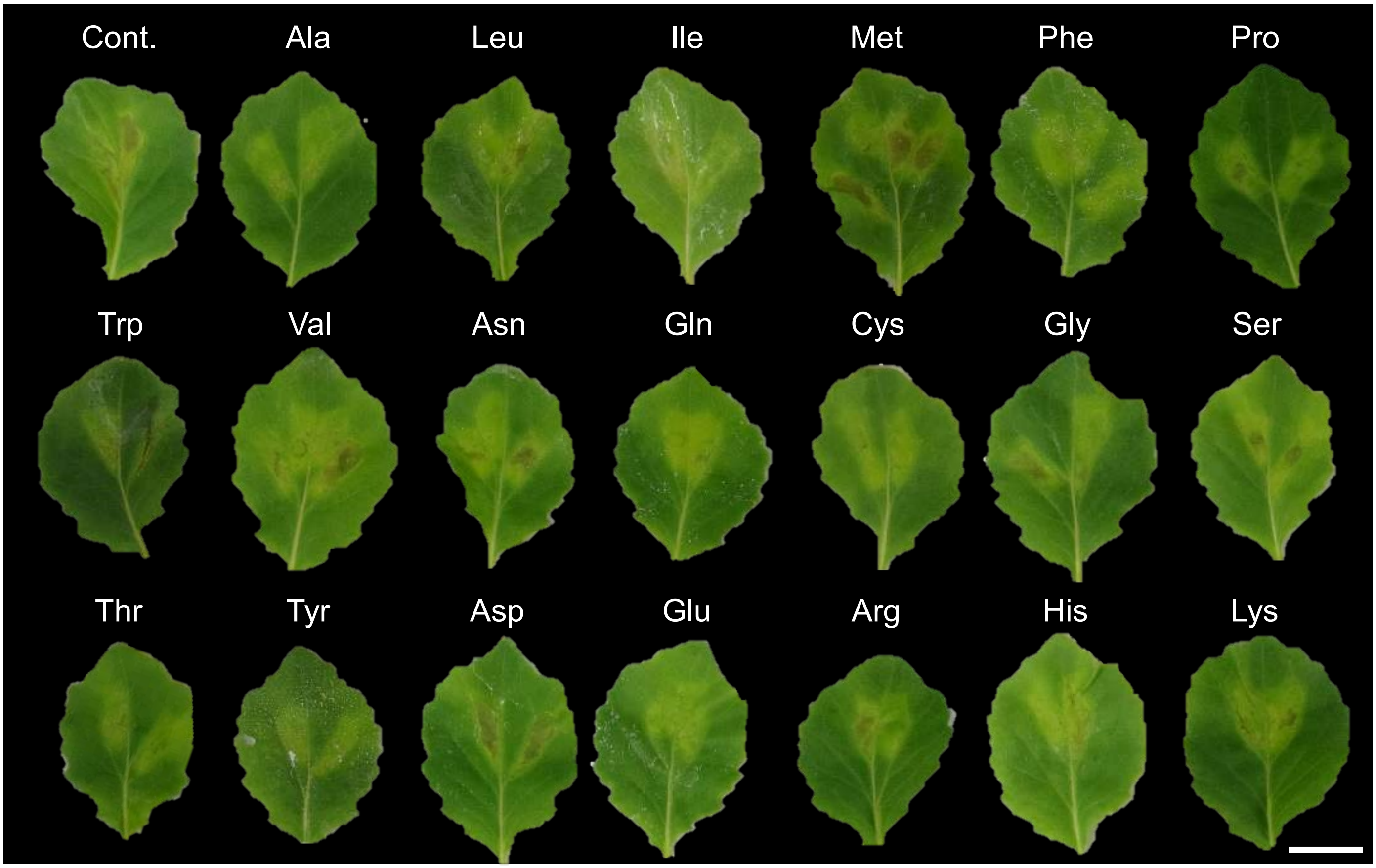


**Supplementary Fig. S3**. **Disease symptoms of amino acids-treated and untreated cabbage leaves after *Pseudomonas cannabina* pv. *alisalnesis* KB211 syringe-inoculation.** Cabbage plants were syringe-inoculated with 5x $10^5$ CFU/ml of inoculum 24 h after amino acids spray-treatment (25 mM), including non-polar amino acids (i.e., Ala, Leu, Ile, Met, Phe, Pro, Trp, and Val), polar amino acids (i.e., Asn, Gln, Cys, Gly, Ser, Thr, and Tyr), acidic amino acids (i.e., Asp and Glu), and alkaline amino acids (i.e., Arg, His, and Lys). Water was used as a control. The leaves were photographed at 5 dpi. Scale bars show 2 cm.

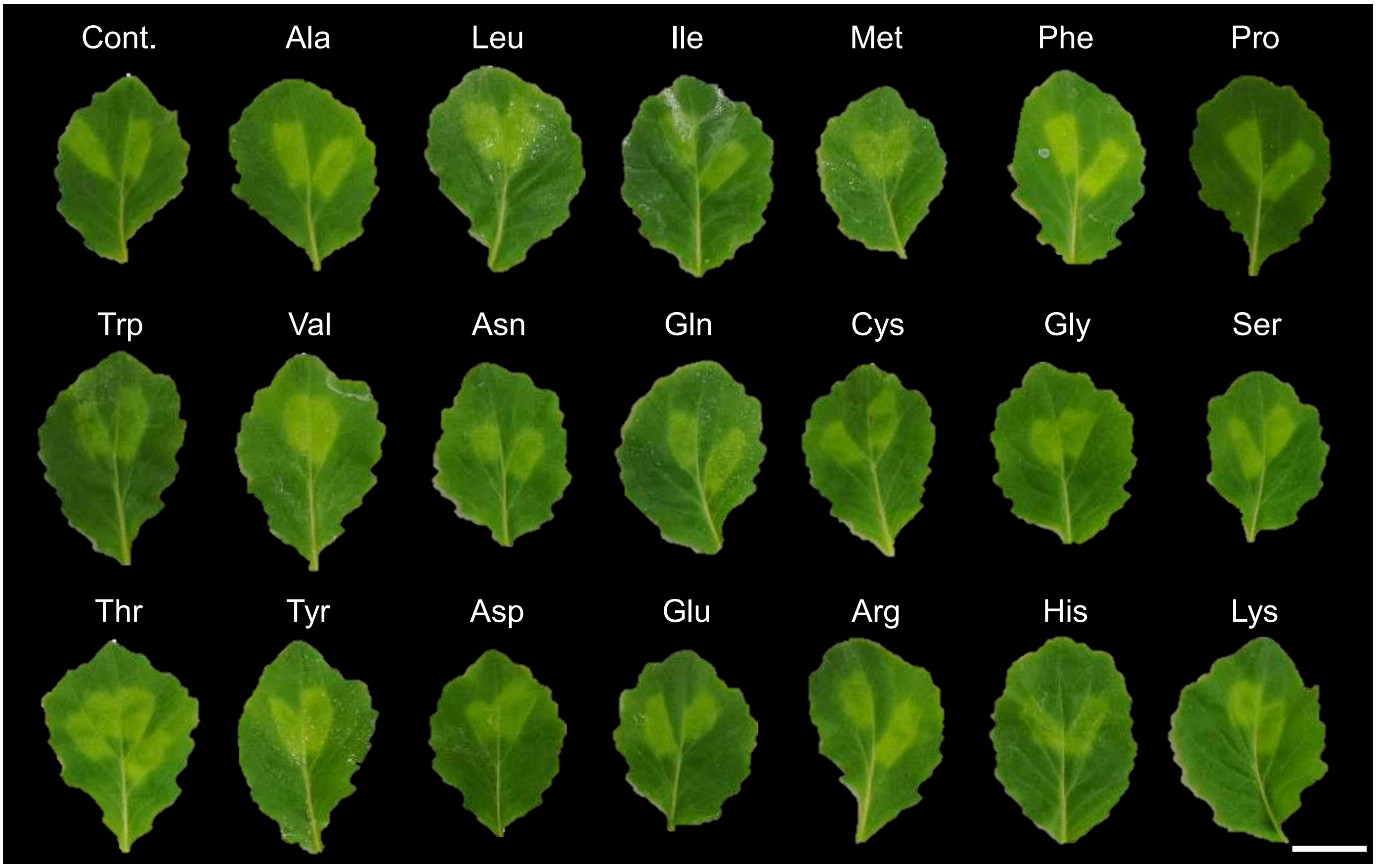


**Supplementary Fig. S4**. **Disease symptoms of amino acids-treated and untreated cabbage leaves after *Pseudomonas cannabina* pv. *alisalnesis* KB211 syringe-inoculation.** Cabbage plants were syringe-inoculated with 5x $10^4$ CFU/ml of inoculum 24 h after amino acids spray-treatment (25 mM), including non-polar amino acids (i.e., Ala, Leu, Ile, Met, Phe, Pro, Trp, and Val), polar amino acids (i.e., Asn, Gln, Cys, Gly, Ser, Thr, and Tyr), acidic amino acids (i.e., Asp and Glu), and alkaline amino acids (i.e., Arg, His, and Lys). Water was used as a control. The leaves were photographed at 5 dpi. Scale bars show 2 cm.

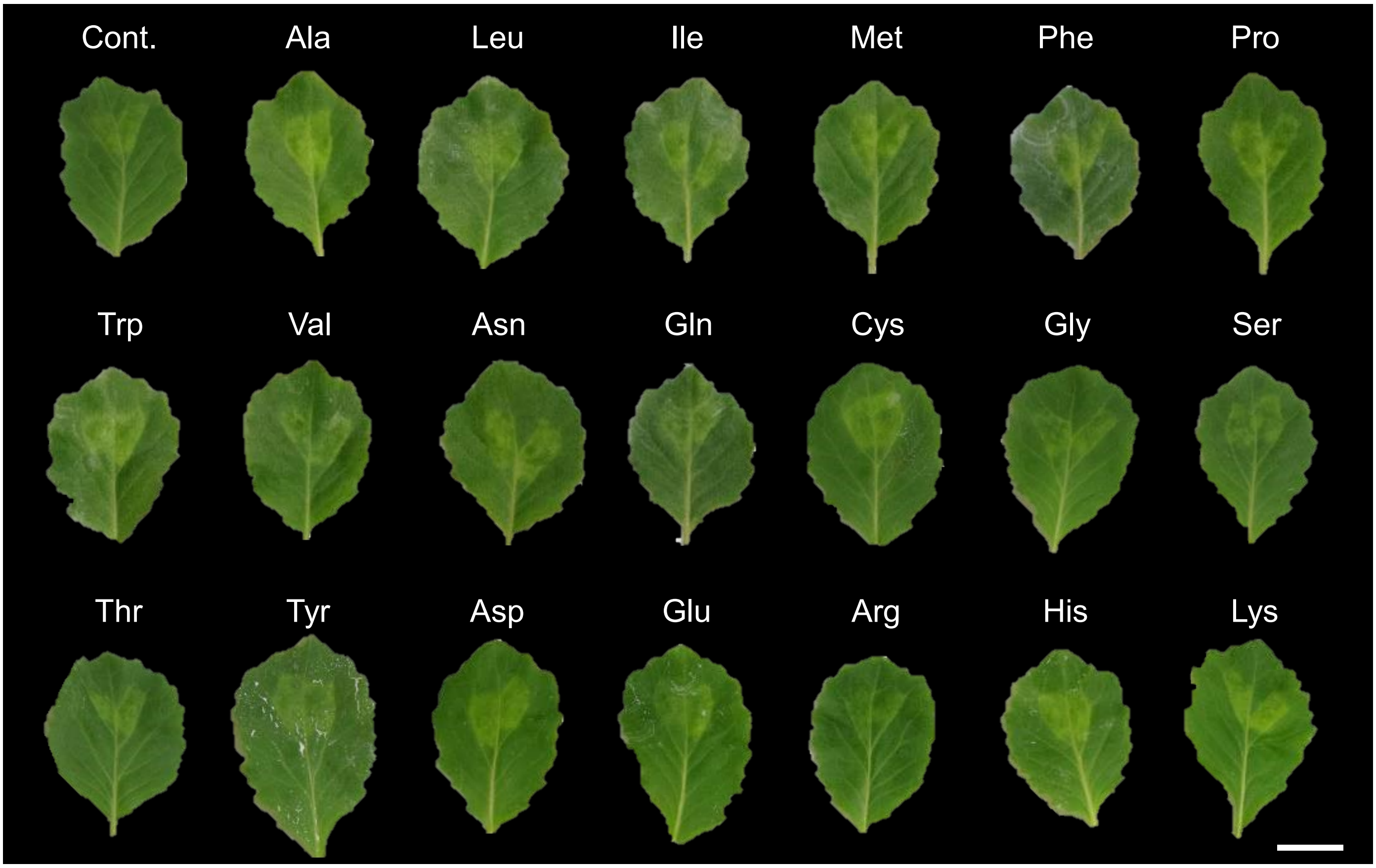


**Supplementary Fig. S5**. **Disease symptoms of amino acids-treated and untreated cabbage leaves after *Pseudomonas cannabina* pv. *alisalnesis* KB211 syringe-inoculation.** Cabbage plants were syringe-inoculated with 5x $10^3$ CFU/ml of inoculum 24 h after amino acids spray-treatment (25 mM), including non-polar amino acids (i.e., Ala, Leu, Ile, Met, Phe, Pro, Trp, and Val), polar amino acids (i.e., Asn, Gln, Cys, Gly, Ser, Thr, and Tyr), acidic amino acids (i.e., Asp and Glu), and alkaline amino acids (i.e., Arg, His, and Lys). Water was used as a control. The leaves were photographed at 5 dpi. Scale bars show 2 cm.

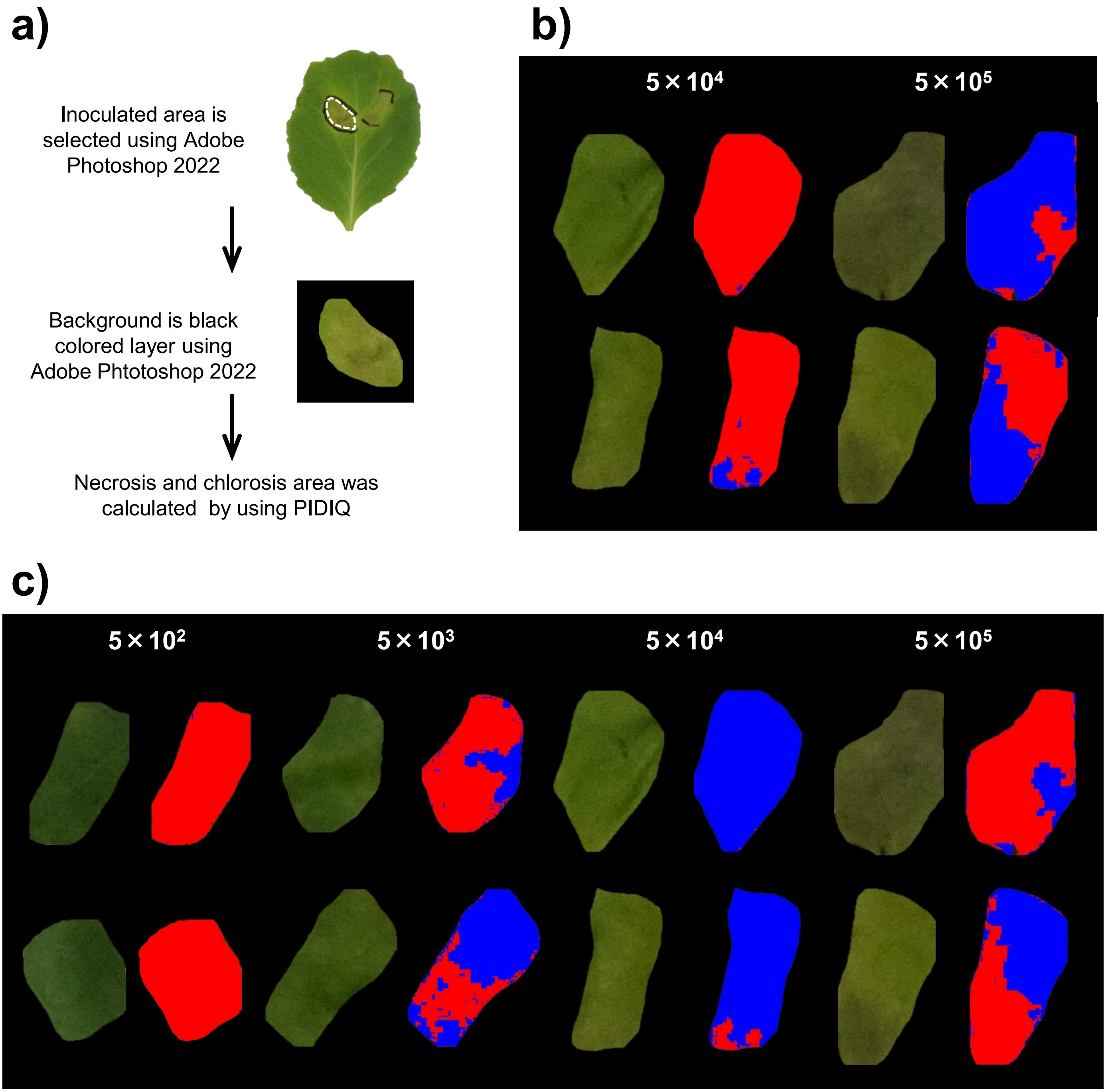


**Supplementary Fig. S6. Necrosis and chlorosis area of cabbage plants after *Pcal* inoculation.** (a) Workflow of the PIDIQ method. The area that has been circled with a white dotted line was selected as an inoculated area by using Adobe Photoshop 2022. (b) Necrosis areas detected by the ImageJ-based PIDIQ method as described in the methods. Necrosis and other areas are colored in blue and red, respectively. (c) Chlorosis areas detected by the ImageJ-based PIDIQ method as described in the methods. Chlorosis and other areas are colored in blue and red, respectively.
